# miR472 deficiency enhances *Arabidopsis thaliana* defence without reducing seed production

**DOI:** 10.1101/2022.12.13.520224

**Authors:** Francois Vasseur, Patricia Baldrich, Tamara Jiménez-Góngora, Luis Villar-Martin, Detlef Weigel, Ignacio Rubio-Somoza

## Abstract

After having co-existed in plant genomes for at least 200 million years, the products of microRNA (miRNA) and Nucleotide-Binding Leucine Rich Repeat protein (NLR) genes formed a regulatory relationship in the common ancestor of modern gymnosperms and angiosperms. From then on, DNA polymorphisms occurring at miRNA target sequences within NLR transcripts must have been compensated by mutations in the corresponding mature miRNA sequence, therefore maintaining that regulatory relationship. The potential evolutionary advantage of such regulation remains largely unknown and might be related to two mutually non-exclusive scenarios: miRNA-dependent regulation of NLR levels might prevent defence mis-activation with negative effects on plant growth and reproduction; or reduction of active miRNA levels in response to pathogen derived molecules (PAMPS and silencing suppressors) might rapidly release otherwise silent NLR transcripts for rapid translation and thereby enhance defence. Here, we used *Arabidopsis thaliana* plants deficient for miR472 function to study the impact of releasing its NLR targets on plant growth and reproduction and on defence against the fungal pathogen *Plectospharaella cucumerina*. We show that miR472 regulation has a dual role, participating both in the tight regulation of plant defence and growth. MIM472 lines, with reduced active miR472, are more resistant to pathogens and, correlatively, have reduced relative growth compared to wild-type plants. However, despite MIM472 lines flower at the same time than their wild-type, the end of their reproductive phase is delayed, and they exhibit higher adult biomass, resulting in similar seed yield as the wild-type. Our study highlights how negative consequences of defence activation might be compensated by changes in phenology and that miR472 reduction is an integral part of plant defence responses.

## Introduction

Pathogens are arguably one of the main threats to an organism’s life. Accordingly, animals and plants have developed sophisticated molecular mechanisms to perceive such threats and defend themselves from invaders. Plants continuously monitor the potential presence of pathogens with dedicated receptors at their cell surface. These proteins, named pattern recognition receptors (PRR), detect molecular signatures conserved in broad classes of microorganisms (DeFalco and Zipfel, 2021). In plants, PRR expression is often cell-type and developmental-stage specific to match the most critical targets of infection (Beck et al., 2014, Emonet et al., 2021). The patterns detected by PRRs refer to molecular signatures produced by pathogens and dubbed pathogen- or microbe-associated molecular patterns (PAMPs/MAMPs). Extracellular recognition of PAMPs by PRRs constitutes a first layer of defence, PAMP-triggered immunity (PTI), which encompasses responses that modify the host’s defensive and developmental status, both at cellular and organismal levels (Bartels and Boller, 2015).

A second surveillance system is dedicated to detecting the intracellular presence and action of pathogen-derived molecules, effectors, used to manipulate the host for the pathogen’s benefit. This second layer of defence relies on proteins from the Nucleotide-binding site leucine-rich repeat superfamily (NLR) (Adachi et al., 2019). NLRs also guard the regular functioning of key host proteins that are hubs in fundamental cellular processes and that are therefore often targets of pathogen intervention. Changes in these host guardees activate NLRs, leading to Effector-Triggered Immunity (ETI) (Jones et al., 2016). While PTI and ETI have initially been considered as independent arms of plant defence, they are now recognized as being closely intertwined and sharing many signalling components, including the participation of hormones such as salicylic (SA) and jasmonic acid (JA) (Lu and Tsuda, 2021).

Inappropriate activation of plant defences often leads to impairment of plant development, growth, and fitness, in both natural settings and in controlled growth conditions (Zust and Agrawal, 2017). Such defence-related trade-offs conform to the assumption that organisms have limited resources to allocate to different physiological processes (Herms and Mattson, 1992). Thus, the increase of resources allocated to defence will result in a proportional decrease in those available for other processes such as growth and/or reproduction. Over the last decades, these trade-offs have been intensively investigated through different approaches in a wide range of organisms (de Jong et al., 200, de Vries et al., 2017, Defossez et al., 2018, Griffiths et al., 2018, Lind et al., 2013, Naseenn et al., 2015). Some studies have supported their existence in plants (Karasov et al., 2014, Tian et al., 2003), although others have failed to detect such trade-offs (Barto and Cipollini, 2005, McGuire and Agrawal, 2005).

To avoid inappropriate activation of defence, timing and magnitude of NLR-based responses are under tight genetic control at the transcriptional, post-transcriptional, translational, and post-translational levels (Cheng et al., 2011, Gloggnitzer et al., 2014, Lai and Eulgem, 2018, Shao et al., 2019, Wu et al., 2017). NLR expression is controlled at the post-transcriptional level by RNA decay and RNA silencing (Gloggnitzer et al., 2014, Shivaprasad et al., 2012, Zhai et al., 2011). RNA silencing in turn relies on two classes of small RNAs (sRNAs); micro RNAs (miRNAs) and small interfering RNAs (siRNAs). miRNAs have emerged as central regulators of plant immunity by directly regulating the expression of NLR genes in a wealth of plant species (Zhang et al., 2016). The impact of miRNA-mediated NLR regulation goes beyond their direct targets, and amplification of the small RNA response serves to regulate additional secondary NLR targets (Zhai et al., 2011). Such amplification starts with conversion of primary targeted transcripts into double-stranded RNAs (dsRNAs) by RNA-dependent RNA Polymerase 6 (RDR6) with 22 nucleotide long miRNAs as trigger. Subsequently, the dsRNAs are recognized and processed by Dicer-Like 4 (DCL4), giving rise to phased secondary siRNAs (phasiRNAs), which can target additional NLR transcripts based on sequence complementarity. Attenuation of miRNA-mediated NLR suppression results in enhanced resistance to viruses, bacteria, oomycetes and fungal pathogens in several plant species, such as Arabidopsis, tomato and barley (Canto-Pastor et al., 2019, Liu et al., 2014, Lopez-Marquez et al., 2021, Shivaprasad et al., 2012).

NLR proteins were already present in the common ancestor of the green lineage (Shao et al., 2019), while miRNAs seem to have evolved independently in chlorophyte green and in brown algae as well as in land plants (Tarver et al., 2015). The appearance of miRNAs targeting NLRs can be traced back to the emergence of gymnosperms, although both miRNAs and NLRs were already present in earlier land plants (Zhang et al., 2016). From that moment on, miRNA-dependent NLR regulation have co-evolved. Different clades of NLR genes themselves evolve at different speeds. Some clades rapidly expand through the generation of NLR paralogs which in turn undergo frequent gene conversion. Other NLRs mainly evolve through presence/absence polymorphisms. Rapidly evolving NLRs, which seem to be more likely to be under miRNA regulation, have been proposed to be particularly beneficial in the presence of new pathogen effectors, widening the resistance spectrum against pathogens (Zhang et al., 2016). Likewise, new miRNA families appear to be continuously generated throughout evolution, and at least eight independent miRNA families having been described as NLR regulators in different groups of plants. Noteworthy, unrelated miRNAs converged to target sequences that encode portions of the highly conserved and functional P-Loop protein domain in their NLR targets. NLRs are under selection from highly dynamic pathogen threats, hence, it has been suggested that NLRs are the driving force of this co-evolution by prompting miRNA mutation and selection to, in turn, maintain their control on NLR expression (Zhang et al., 2016).

Thus, two scenarios are possible. On the one hand, miRNAs have been evolutionary recruited by the NLR networks to reduce collateral damage from inappropriate NLR activation. On the other hand, miRNAs were added as a third layer to the pathogen response, with miRNAs acting as indirect pathogen sensors.

We reasoned that if the role of NLR-regulating miRNAs is primarily to prevent undesired developmental or physiological effects, plants lacking these miRNAs should also be compromised in growth and reproduction. Alternatively, limited growth/reproduction defects would favour a scenario in which miRNAs have a direct role in regulating defence.

The A. *thaliana* reference genome encodes two unrelated miRNAs that regulate NLR expression, miR472 and miR825-5p (Boccara et al., 2014, Lopez-Marquez et al., 2021). miR472 regulates a group of genes belonging to the coiled-coil-NLR (CC-NLR) sub-class, while miR825-5p primarily targets members from the Toll-interleukin-receptor-like-NLR (TIR-NLR) subfamily. Deficient miR472-mediated NLR regulation increases defence response against the bacterial pathogen *Pseudomonas syringae* (Boccara et al., 2014).

Here, we have characterized the role of *A. thaliana* miR472-dependent NLR regulation in plant defence against the fungal pathogen *Plectospharaella cucumerina* and its involvement in different trade-offs. We report that miR472 downregulation is part of the response against pathogens triggered by the detection of PAMPs produced by the hemibiotrophic fungus *Plectosphaerella cucumerina*. Plants deficient in miR472 have lower rosette relative growth rate (RGR) in the absence of pathogens, consistent with a growth-defence trade-off. However, miR472 deficiency is also associated with a longer life cycle, leading to bigger size at reproduction, and thus without a net effect on overall seed production.

## Results

### Fungal PAMPs reduce miR472 levels

Treatments with the bacterial PAMP flg22 downregulates *A. thaliana* miR472 (Boccara et al., 2014, Su et al., 2018). Since we have also described a similar trend for some plant miRNAs as part of fungal elicitor-triggered PTI, including the NLR-regulating miR825-5p (Lopez-Marquez et al., 2021, Salvador-Guirao et al., 2018), we tested whether miR472 was also responsive to fungal PAMPs. To that end, we measured the expression of miR472 and its NLR target *At5g43740* after elicitor treatment. Four-week old wild-type Col-0 plants were treated with either *P. cucumerina*-derived elicitors or with a mock solution, and samples were collected at different time points. miR472 downregulation was readily detected as early as 30 minutes after elicitor treatment (Fig. 1A). Expressoin at later time points was variable, but always lower than in control plants. Such a fluctuating expression pattern resembles the one previously reported in response to flg22 (Boccara et al., 2014). The expression of At5g43740, one of the miR472 target genes, did not increase until 60 minutes after elicitor treatment, and its expression stayed elevated at later time points (Fig. 1B). These results are consistent with miR472 acting through its target At5g43740 in the response to fungal elicitors.

**Figure 1.**
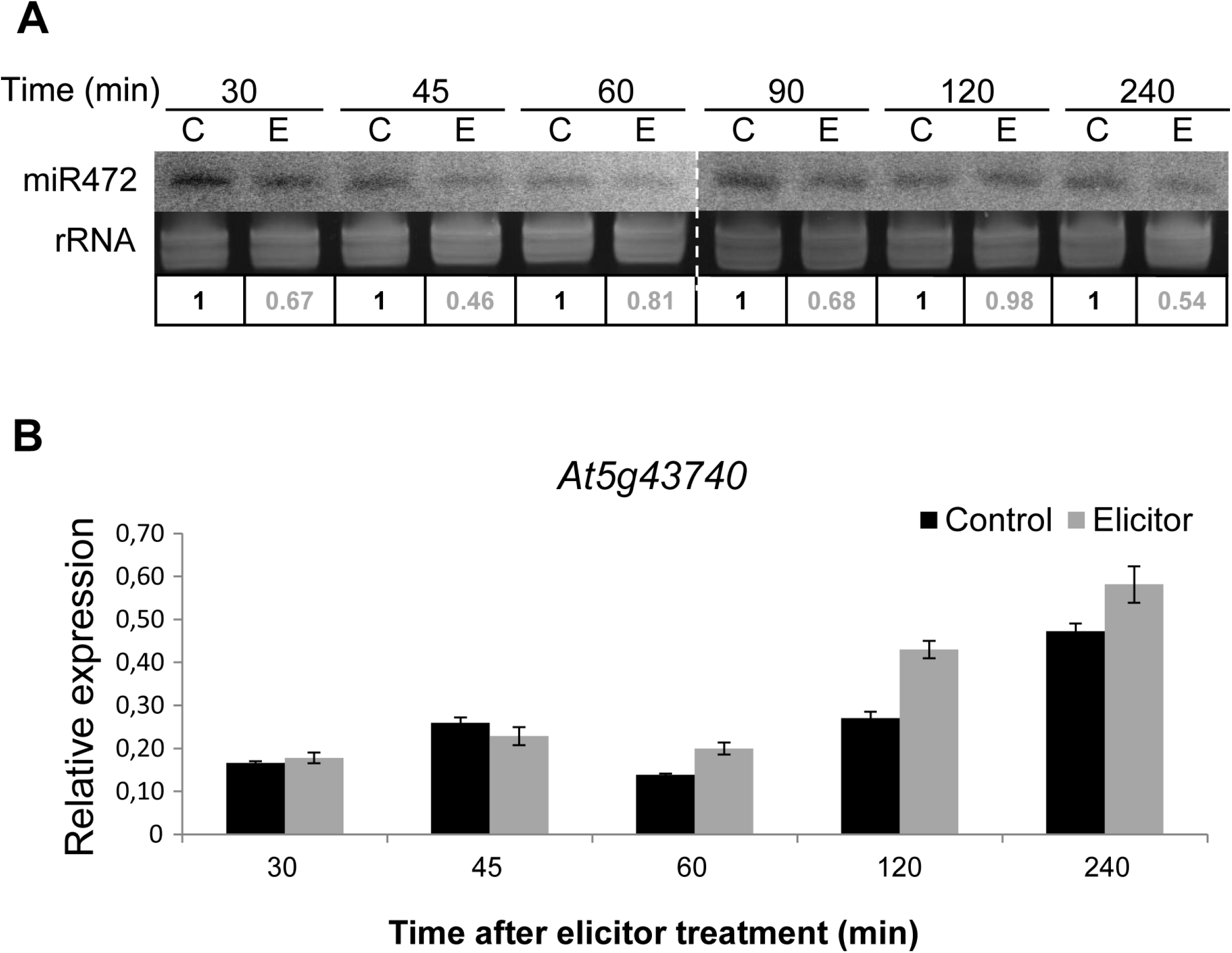
Fungal elicitors trigger a downregulation of miR472 and up-regulation of its target in *Arabidopsis thaliana*. **A**. Accumulation of miR472 determined by small RNA Northern blot in a time course of wild-type Col-0 plants treated with *P. cucumerina* elicitors (300 μg/ml), or mock solution (control). Northern blots were hybridized with a *γ32P-ATP* end-labelled probe complementary to miR472. Ribosomal RNA (rRNA) was used as a loading control. Relative band density was determined using ImageJ and normalized to the intensity of its corresponding rRNA control and, subsequently, to the corresponding control time point. **B**. Relative expression of miR472 target CC-NBS-LRR (At5g43740), as determined by reverse transcription quantitative PCR (RT-qPCR). The β-tubulin2 gene (At5g05620) was used as housekeeping for transcript expression normalization. The experiment was repeated for a total of three biological replicates. All replicates presented similar results. The histogram shows results of one of the three biological replicates.

### miRNA472 downregulation increases resistance to Plectosphaerella cucumerina

*Arabidopsis thaliana* plants deficient in miR472 can cope better with bacterial infections than wild-type plants (Boccara, Sarazin et al. 2014). To assess whether miR472 deficiency also contributes to increased resistance against fungal pathogens, we challenged miR472 target mimic (MIM472) plants with reduced miR472 activity (Todesco, Rubio-Somoza et al. 2010) with *P. cucumerina* (Petriacq, Stassen et al. 2016). MIM472 expression results in miR472 degradation and in higher levels of expression of at least one of its NLR targets, At5g43740 (Fig. 2A, Fig. S1). Four-week old MIM472 and control plants were inoculated with fungal spores and plant survival was scored 12 days-post-inoculation (dpi). MIM472 plants outperformed both wild-type and empty vector plants, with up to 70% survival, compared to only around 30% of surviving control plants (Fig. 2B, C).

**Figure 2.**
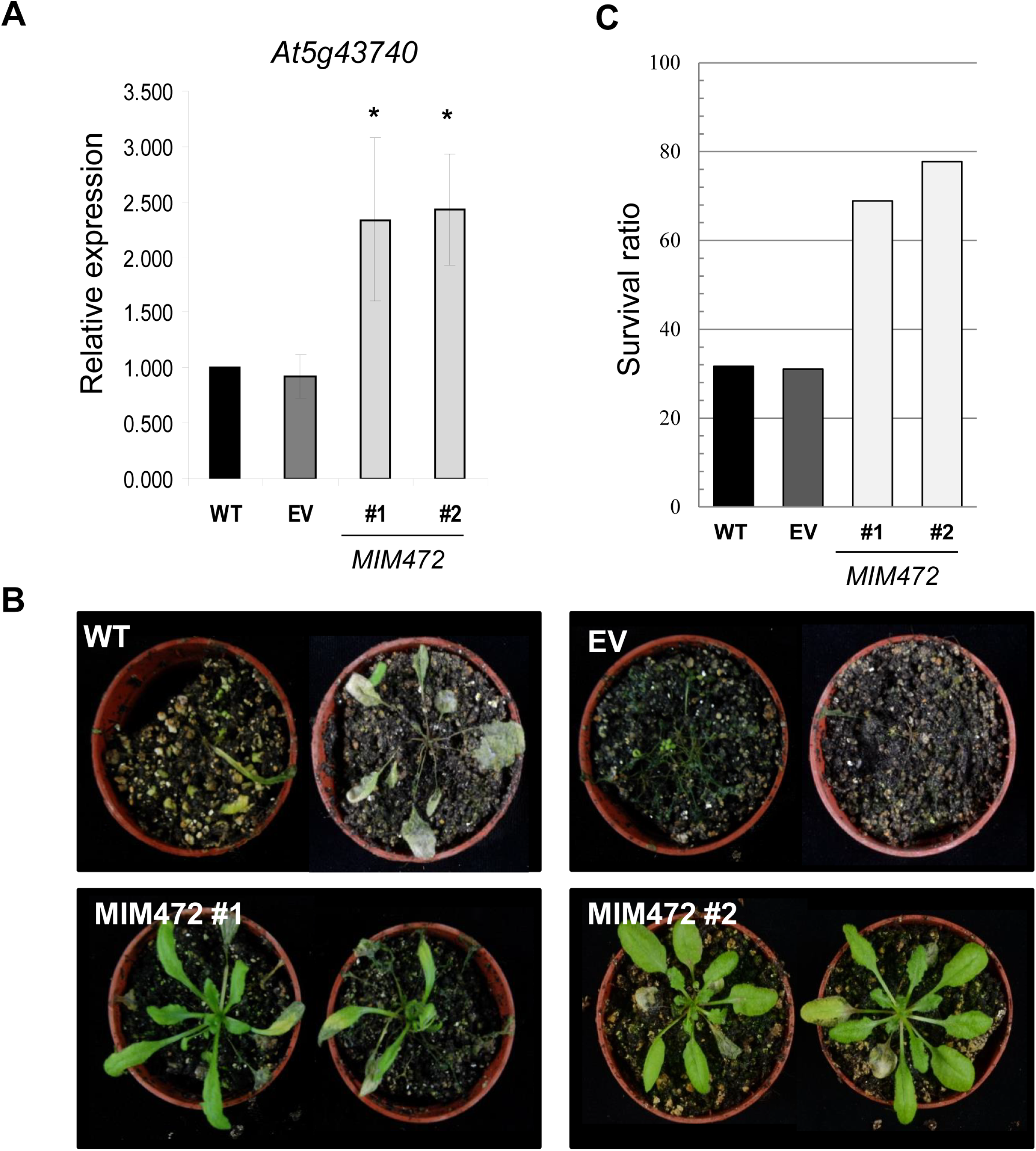
MIM472 plants are more resistant to *P. cucumerina* infection. **A**. Relative expression of a miR472 CC-NBS-LRR target (At5g43740), as determined by RT-qPCR. The β-tubulin2 gene (At5g05620) was used as a housekeeping gene for transcript expression normalization. Three biological and technical replicates were included in the assay. Significance codes (ANOVA-test): *: *p<* 0.05. **B**. Plants 12 days after spray-inoculation with *P. cucumerina* spores (1.10^6^ spores/ml). Six biological replicates were used for each genotype. All biological replicates behaved similarly. Histogram shows results from one of the three biological replicates. **C**. Survival ratio of plants infected with *P. cucumerina* spores (1.10^6^ spores/ml) at 12 days post infection. Six biological replicates were used for each genotype. All technical replicates behaved similarly, showing results of one of the three technical replicates.

Prior to pathogen challenge, MIM472 plants did not show reduced photosynthetic capacity, often a hallmark of defence responses (Berger et al., 2007, Rousseau et al., 2013) (Fig. S2). Additionally,theseMIM472 plants did not present a correlation between increased resistance (Fig. 2B, C) and the activation of hormone-mediated defence programs, as inferred from the expression levels of marker genes for either JA- (*PDF1*.*2, VSP2*) or SA-dependent (PR1, PAD4) pathways (Fig. S3). Together, these results show that miR472 regulates NLR targets involved in anti-fungal defence and that its deficiency does not constitutively activate defence markers.

### Impaired *miRNA 472 activity decreases rosette relative growth rate* (RGR)

To evaluate a possible contribution from miR472 to NLR-dependent defence and growth/fitness trade-offs, we monitored development, growth, and reproductive trajectories in MIM472 and wild-type plants. Using the Raspberry Pi Automated Phenotyping Array (RAPA) platform (Vasseur, Bresson et al. 2018) (Fig. 3A), we tracked rosette area in plants from two MIM472 lines during ontogeny and until plant maturity (i.e., when fruits are ripening, which is considered as the end of reproduction). Sigmoid growth curves were fitted on every individual to estimate growth parameters (Fig. 3B). Rosette relative growth rate (RGR) was estimated as the rate of increase in rosette area per unit existing area (mm^2^ d^-1^ cm^-2^) at the inflection point of the growth curve (i.e. when absolute growth rate was maximal). The two MIM472 lines had significantly lower RGR than Col-0 wild-type plants (Kruskal-Wallis P < 0.05; Fig. 4C), but RGR reduction was not associated with a decrease of chlorophyll fluorescence, as inferred from the *F*_v_/*F*_m_ ratio measured on three-week old, unchallenged plants (Fig. S2). This ratio reflects the efficiency of the photosystem to transport electrons (Bresson et al., 2015, Murchie and Lawson, 2013), and it is often reduced with the onset of necrosis following pathogen attack (Berger et al., 2007, Rousseau et al., 2013). Overall, these results point to a growth-defence trade-off in plants with reduced miR472 activity. Moreover, the reduction of growth seems to mainly result from an effect of miR472 on resource allocation rather than resource acquisition through photosynthesis.

**Figure 3.**
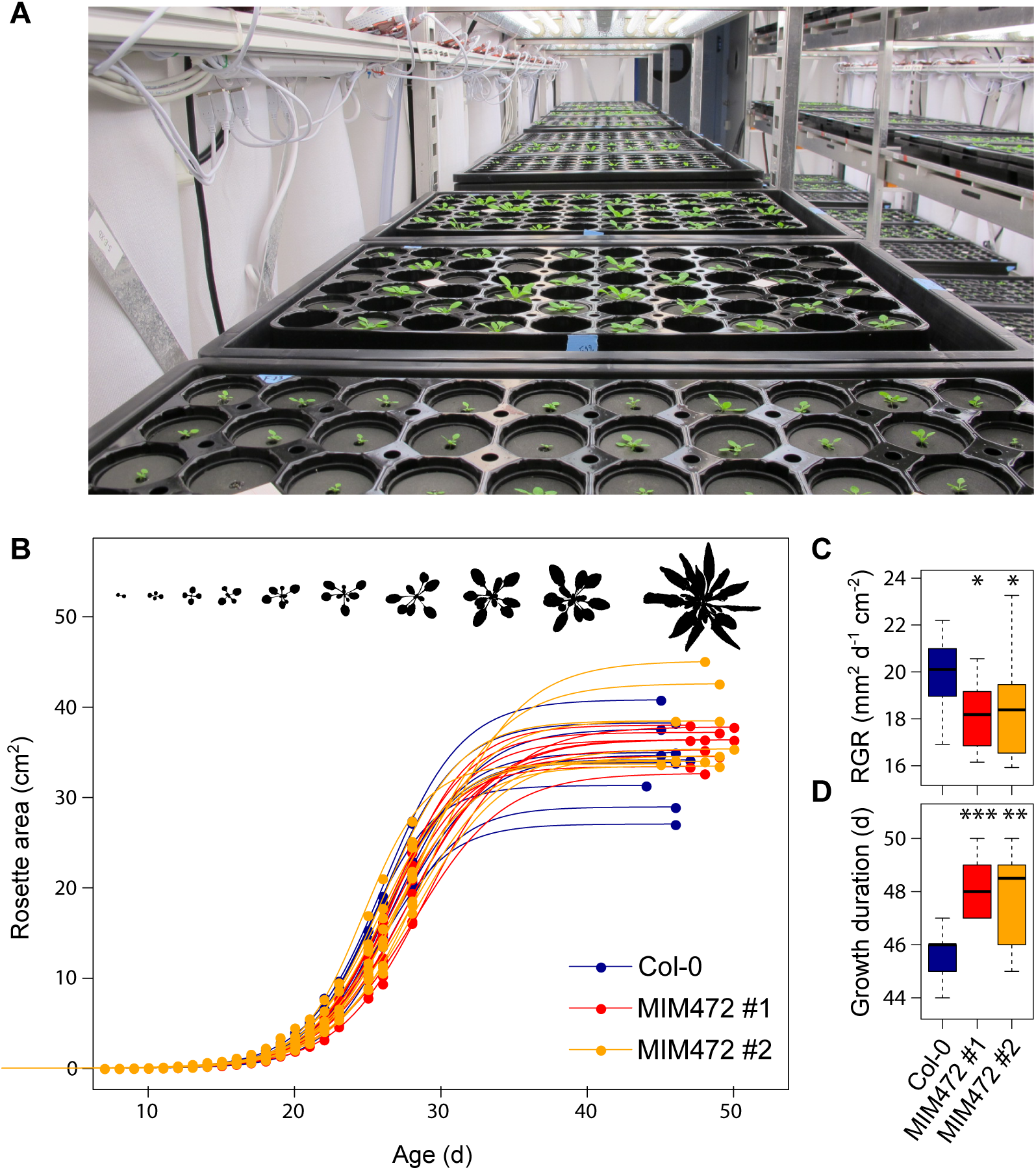
Variation of growth and phenology in MIM472 lines. **A**. Photography of the Raspberry Pi Automated Phenotyping Array (RAPA) phenotyping platform, with trays of *Arabidopsis thaliana* grown in highly controlled conditions **B**. Growth curves of two MIM472 lines (MIM472 #1 in red and MIM472 #2 in orange) and the wild-type Col-0 (in blue) grown in RAPA (*n* = 10). Dots are the rosette area (cm^2^) measured from image analysis, with the last ones measured at maturity when plants were harvested. Curves were fitted with logistic models. At the top of the graph is a representative rosette (Col-0) with images taken during ontogeny. **C**. Differences in relative growth rate (RGR) between the two MIM472 lines and the wild-type Col-0 (*n* = 10). **D**. Differences in age at bolting between the two MIM472 lines and the wild-type Col-0 (*n* = 10). (C) Differences in age at maturity between the two MIM472 lines and the wild-type Col-0 (*n* = 10). Significance codes (Kruskal-Wallis non-parametric tests): ***: *p* < 0.001; **: *p* < 0.01; *: *p* < 0.05; NS: *p* > 0.05.

**Figure 4.**
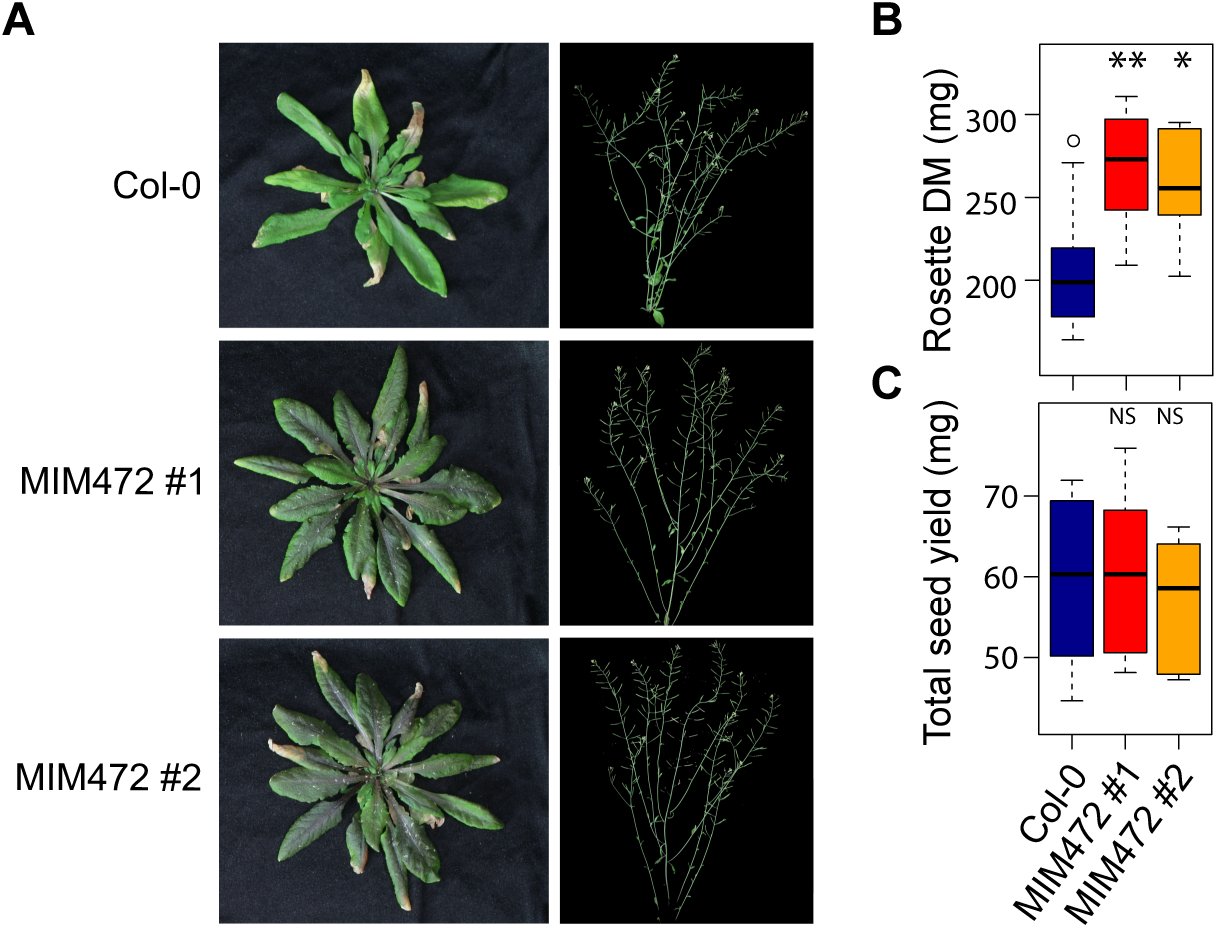
Variation of rosette dry mass and seed yield in MIM472 lines. **A**. Examples of pictures taken at maturity (end of reproduction). **B**. Differences in rosette dry mass (DM, mg) between the two MIM472 lines and the wild-type Col-0 (*n* = 10). **C**. Differences in total seed yield (mg) between the two MIM472 lines and the wild-type Col-0 (*n* = 10). Significance codes (Kruskal-Wallis non-parametric tests): ***: *p* < 0.001; **: *p* < 0.01; *: *p* < 0.05; NS: *p* > 0.05.

### Growth and phenological adjustments in MIM472 are associated with fitness homeostasis

Despite significant differences in RGR, the timing of reproduction, measured as the age at bolting, was similar between MIM472 lines and the wild-type Col-0 (Kruskal-Wallis P > 0.05 for MIM472-1, and P = 0.02 for MIM472-2; Fig. S3A). In contrast, we noticed that MIM472 individuals ended their reproduction two days later, on average, than Col-0 (Kruskal-Wallis P < 0.01; Fig. 3D). Accordingly, MIM472 lines had longer reproductive phase than their wild-type (Kruskal-Wallis P < 0.01 for MIM472-1, and P = 0.11 for MIM472-2; Fig. S3B). This prompted us to test whether the coordinated changes in RGR and reproductive growth duration could buffer the effects of defence-related processes on fitness in the MIM472 lines by determining rosette dry mass at maturity (i.e., end of reproduction) as well as total seed yield (Fig. 4). Both MIM472 lines had significantly higher rosette dry mass at maturity compared to Col-0 (Kruskal-Wallis P < 0.05; Fig. 4B). By contrast, the seed yields of MIM472 and Col-0 wild-type plants were not significantly different (Kruskal-Wallis P > 0.05; Fig. 4C), averaging at 60 mg of seeds per plant for all genotypes, indicating that an extended growth phase compensates for slower growth rate.

## Discussion

It has been speculated that miRNA play an essential role to reduce the negative impact of NLR expression on plant development (Gonzalez, Muller et al. 2015, Canto-Pastor, Santos et al. 2019). Here, we characterized the phenotypic costs of higher NLR activity in mutants impaired in miR472-dependent regulation. Our results are consistent with miRNA-dependent regulation of defence is associated with few fitness costs in *Arabidopsis thaliana*. Likewise, our results suggest that an advantage of miRNA-mediated NLR regulation allows for cell host reprogramming upon detection of pathogen threats. Thus, pathogen-derived dysfunction of host-proteins and sRNAs triggers NLR-mediated defence.

It is generally assumed that the metabolic cost of defence activation should be associated with a concomitant decrease of growth and reproductive output (Herms and Mattson, 1992, Karasov et al., 2017, Obeso, 2002). Indeed, resource allocation to growth and reproduction are expected to be both negatively, and similarly, impacted by an increase in defence metabolism. However, our results suggest that the relationships between growth, reproduction, and defence are complex. We found that higher defence in MIM472 lines is associated with slower relative growth rate but no reduction in seed production. Further analysis showed that the slow-growing MIM472 lines have a longer life cycle, and specifically a longer reproductive phase. Negative relationships between growth rate and lifespan are commonly reported in large comparative studies among *A. thaliana* ecotypes (Debieu et al., 2013, Sartori et al., 2019, Vasseur et al., 2014, Vasseur et al., 2018a), as well as among plant species (Reich, 2014, Wright et al., 2004)). This is generally interpreted as the result of physiological constraints, such as the necessity to allocate resources to structural compounds to increase mechanical resistance and support a long lifespan. In *A. thaliana*, delayed reproduction is associated with a higher accumulation of vegetative biomass. This increases the availability of resources, such as nitrogen, to be possibly remobilized during senescence from the vegetative to the reproductive parts (Killingbeck, 1996, Sartori et al., 2022). Consistently, we found here that up-regulating defence can have a cost on relative growth rate, as hypothesized by physiological considerations about the growth-defence trade-off, while maintaining seed production. We argue that this maintenance of seed production is reached by accumulating more vegetative biomass and extending reproduction to optimise resource resorption from leaves to fruits. Together, our findings suggest that fitness can be maintained upon defence activation through coordinated changes in plant physiology and phenology.

Life history transitions such as the onset of flowering in annual plants are critical steps, tightly regulated at the molecular level to integrate multiple environmental cues (Mouradov et al., 2002). In *A. thaliana*, mechanisms that accelerate reproduction are expected to be selected under water-limited conditions, for instance in Mediterranean regions where the seasonal window with favourable water conditions is short (Vasseur et al., 2018b). Our findings suggest that miRNA activity impacts plant development and phenology, which likely interferes with the metabolic regulation triggered by environment signals such as temperature and day length. Hence, if there is a fitness effect of defence activation in MIM472 lines, it is not due to resource limitation for seed production, but rather on a potential mismatch between environment and development. For instance, an inappropriate activation of defence might delay reproduction and increase the risk of drought in Mediterranean climate. It is likely that the physiological changes induced by the miRNA system of defence regulation might depend on the number of NLR genes targeted and downregulated. Recent work has shown that miRNAs mutate to keep on track with mutations in their targets (Zhang et al., 2016). We propose here that this is not to prevent undesired effects on fitness, but rather to link their upregulation to a failure of the silencing machinery as consequence of the presence of pathogens, and thus, trigger a fast defence reprogramming to cope with those pathogen threats. In line with that, reduction in functional miR472 and miR825-5p as consequence of PAMP detection (this work, Boccara et al., 2014, Lopez-Marquez et al., 2021) and/or the action of silencing suppressors impinging sRNA function, which constitutes a conserved strategy employed by different pathogens during infection (Csorba et al., 2015, Navarro et al., 2008, Qiao et al., 2013, Zhu et al., 2022), are translated into enhanced NLR expression and defence.

Moreover, another reason for explaining conflicting findings about growth- and fitness-defence trade-offs relies on the fact that most plant pathogens target vegetative tissues for infection rather than reproductive organs. For instance, miR472, like miR825-5p, is preferentially expressed in Arabidopsis leaves when compared to inflorescences (Vazquez et al., 2008), which suggest that tissue and/or developmental stage restriction of miRNA-dependent regulation of NLRs would aid to maximise defence response upon pathogen-triggered sRNA dysfunction while minimising its effect on fitness. Further studies, for instance with reciprocal transplants, are necessary to determine to what extent defence regulation by miRNAs might have an environment-dependent cost on fitness, due to changes in life history rather than changes in reproductive output.

## Conclusions

Our study suggests that miRNAs targeting NLRs work to a large extent as sensors for the presence of pathogens. sRNA-mediated NLR expression through miRNAs and amplifying siRNAs render this system sensitive to silencing suppressors targeting many steps in both regulatory pathways, such as those involved in miRNA biogenesis (DCL1, HYL1), miRNA action (AGO1), and in siRNA amplification (RDR6, SGS3, DLC4, DRB4). Our results show that the negative effect on growth from defence activation in MIM472 lines can be attenuated by phenological adjustments leading to higher biomass and fitness homeostasis. Overall, this suggests that growth-defence and fitness-defence trade-offs can be uncoupled due to compensatory mechanisms involving different traits at different ontogenic stages.

## Materials and Methods

### Plant material

MIM472 plants (Col-0 background) were produced and described in (Todesco, Rubio-Somoza et al. 2010). For infection assays and elicitor treatments, Col-0 and MIM472 plants were grown directly on a mixture of soil:perlite:vermiculine (2:1:1) under neutral day photoperiod (12h of light and 12h of dark), at 22C during the day and 20C during the dark, and 60% relative humidity, in controlled environments for four weeks for pathogen assays and until the end of their life cycle for phenotyping assays.

### Infection assays and elicitor treatment

*Plectosphaerella cucumerina* was grown at room temperature in petri dishes containing PDA media (Potato Dextrose Agar, BD Difco), supplemented with chloramphenicol, for 3 weeks. Spores were collected by adding sterile water to the surface of the plate, rubbing with a glass cell spreader and collecting spores released in the water. Spore concentration was adjusted to 1.10^6^ using a Burker counting chamber and a light microscope. Spores were used to spray inoculate six four weeks-old plants/genotype. The progression of the infection was monitored up to 12 days after inoculation. For elicitor treatment, *Arabidopsis thaliana* Col-0 plants were sprayed with an elicitor suspension obtained from *P. cucumerina* (300 μg/ml) as previously described (Coca and San Segundo 2010).

### Expression assays

Total RNA was isolated using TRIzol reagent (Invitrogen). First-strand cDNA was synthesised from 1 ug of TURBO DNAse I (Ambion) or DNAse I (Thermo Fisher) treated total RNA using SuperScript III kit (Invitrogen) or Revertaid First strand kit (Thermo Fisher). Reverse transcription quantitative PCR (RT-qPCR) was performed in a Light Cycler 480, using SYBR green (Roche). Primers were designed using Oligo Analyzer software (Integrated DNA Technologies).

For RT-qPCR, we used b-tubulin2 gene (At5g05620) as a housekeeping for transcript expression normalisation. ΔCt method was used to analyse the results.

For miRNA expression, mature miR472 accumulation was determined by stem-loop RT-qPCR, as described in (Varkonyi-Gasic, Wu et al. 2007). We confirmed the specific amplification of the miR472 using amplicon sequencing.

For Northern blot assays, 20ug or total RNA were fractioned in a 17.5% polyacrylamide denaturing gels containing 8 M urea and transferred to nylon membranes. Probes were design to be complementary to miR472 and end-labelled with γ32P-ATP.

All primers and probes used in this study are indicated in supplemental table 1.

### Phenotyping of growth- and fitness-related traits

Seeds of Col-0, MIM472 #1 and MIM472 #2 were sown in individual pots (*n* = 10), randomly distributed in trays of 30 pots each. Circular pots of 4.6 cm (diameter) x 5 cm (depth) (Pöppelmann, Lohne, Germany) were filled with soil (CL T Topferde; www.einheitserde.de). After germination, plants were thinned to one individual per pot, and trays were moved to the Raspberry Pi Automated Plant Analysis (RAPA) facility (Vasseur et al., Plant Methods 2018), set to 16°C, air humidity at 65%, and 12 h day length, with a PPFD of 125 to 175 µmol m-2 s-1 provided by a 1:1 mixture of Cool White and Gro-Lux Wide Spectrum fluorescent lights (Luxline plus F36W/840, Sylvania, Germany). Trays were randomly positioned in the room and watered every 2 to 4 days.

Growth-related traits were measured using methodologies previously published (Vasseur, Bresson et al. 2018). The RAPA system was used for rosette imaging during the first 28 days of growth. Rosette area (cm^2^) was measured from pictures with imageJ (Schneider, Rasband et al. 2012). Sigmoid growth curves were fitted on all individuals as previously described (Vasseur, Bresson et al. 2018). From the parameters of the fitted functions, we estimated RGR (rosette growth rate at time *t* divided by rosette area at time *t*, mm2 d-1 cm-2) over time. We used RGR at the inflection point of the growth curve from each individual for comparison, which corresponds to the transition point between the vegetative and the reproductive phases, when the absolute growth rate is maximal. We used the inflection point as estimates of bolting stage (in days). We estimated the efficiency of photosynthesis with images of chlorophyll fluorescence, measured with *F*_v_/*F*_m_ ratio at the whole-plant level, after 20 min of dark-adaptation on three-week old rosettes. Chlorophyll fluorescence images were taken with a high-throughput imaging system (Imaging-PAM M-Series, Maxi-version, Heinz Walz GmbH, Germany), and analysed with ImageJ (Schneider, Rasband et al. 2012) to estimate mean *F*_v_/*F*_m_ of each rosette individually (Bresson, Vasseur et al. 2015) for details about chlorophyll fluorescence measurements).

Plants were harvested at maturity, when fruits were ripening. Age at maturity (d) was measured as the duration between germination and the end of reproduction. Rosettes were separated from reproductive parts, photographed with a high-resolution, 16.6 megapixel SLR camera (Canon EOS-1, Canon Inc., Japan), then dried at 65° C for three days and weighed to measure rosette dry mass (DM, mg). Inflorescences were dried at 35 °C for one week in paper bags. Seeds were collected and sifted to remove shoot fragments and weighed.

### Statistical analyses

Statistical significance from expression assays (RT-qPCR) was determined using an ANOVA test adjusted (p<0,05).

For phenotypic analysis, logistic functions were fitted with *nls* in R. Differences in phenotypic traits between Col-0 and the two MIM472 lines were tested with non-parametric Kruskal-Wallis tests. All statistical analyses were performed in R v3.2.3.

## Supporting information

SOM Fig 1

SOM Fig 2

SOM Fig 3

SOM Table 1

## Acknowledgements

We would like to thank Miguel Angel Blazquez, Pere Puigdomenech and members of the MoRE Lab for critical reading of the manuscript. Work at the Max Planck Institute was funded by the Max Planck Society (D.W.). The work in the MoRE laboratory is funded by BFU-2014-58361-JIN (funded by MCIN/AEI/ 10.13039/501100011033 and by “ESF Investing in your future”), RYC-2015-19154 (funded by MCIN/AEI/ 10.13039/501100011033 and by “ESF Investing in your future”), RTI2018-097262-B-I00 (funded by MCIN/AEI/ 10.13039/501100011033 and by “ERDF A way of making Europe”); and through the “Severo Ochoa Programme for Centres of Excellence in R&D” 2016-2019 (SEV-2015-0533) and 2020-2023 (CEX2019-000902-S) funded by MCIN/AEI/ 10.13039/501100011033 and the CERCA programme from the Generalitat de Catalunya. L. V-M was supported by BES-2016-076986 (funded by MCIN/AEI/ 10.13039/501100011033 and by “ESF Investing in your future”). Tamara Jiménez-Góngora was recipient of a Postdoctoral Fellowship from the CRAG through the “Severo Ochoa Programme for Centres of Excellence in R&D” 2016-2019 (SEV-2015-0533).

## Figure legends

**Supplemental Figure 1. MIM472 plants show lower levels of mature miR472**.

Validation of lower levels of mature miR472 in MIM472 plants compared to WT by RT-qPCR. As described in (Todesco, Rubio-Somoza et al. 2010) functional MIM constructs trigger degradation of the decoyed miRNA resulting in lower levels of its mature form. Two biological and technical replicates were included in RT-qPCR assays. Significance codes (ANOVA-test): *: *p<* 0.05.

**Supplemental Figure 2. Variation of photosynthesis efficiency in MIM472 lines**.

Photosynthesis efficiency was measured with chlorophyll fluorescence (Fv/Fm ratio, unitless) on three-week old plants (*n* = 10). Pictures used as examples here were coloured with imageJ. Significance codes (Kruskal-Wallis non-parametric tests): ***: *p* < 0.001; **: *p* < 0.01; *: *p* < 0.05; NS: *p* > 0.05.

**Supplemental Figure 3. Expression pattern of defence-related hormone marker genes**.

**A**. Expression analysis of the SA marker gene PR1 by RT-qPCR in MIM472 and WT plants.

**B**. Expression analysis of the SA marker gene PAD4 by RT-qPCR in MIM472 and WT plants.

**C**. Expression analysis of the JA marker gene PDF1.2 by RT-qPCR in MIM472 and WT plants.

**D**. Expression analysis of the JA marker gene VSP2 by RT-qPCR in MIM472 and WT plants.

Two biological and technical replicates were included in RT-qPCR assays. Significance codes (ANOVA-test): *: *p<* 0.05.

