## Supplementary figures and images for "miR472 deficiency enhances *Arabidopsis thaliana* defence without reducing seed production"

### SOM Fig 1

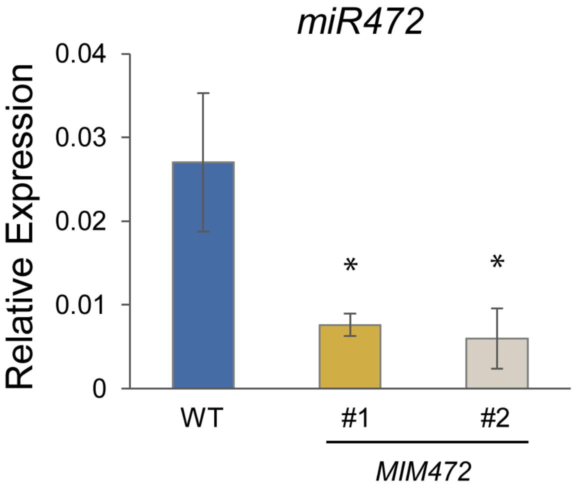

### SOM Fig 2

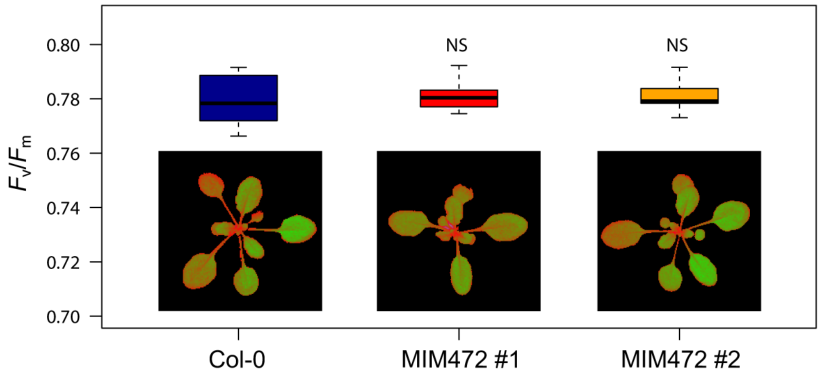

### SOM Fig 3

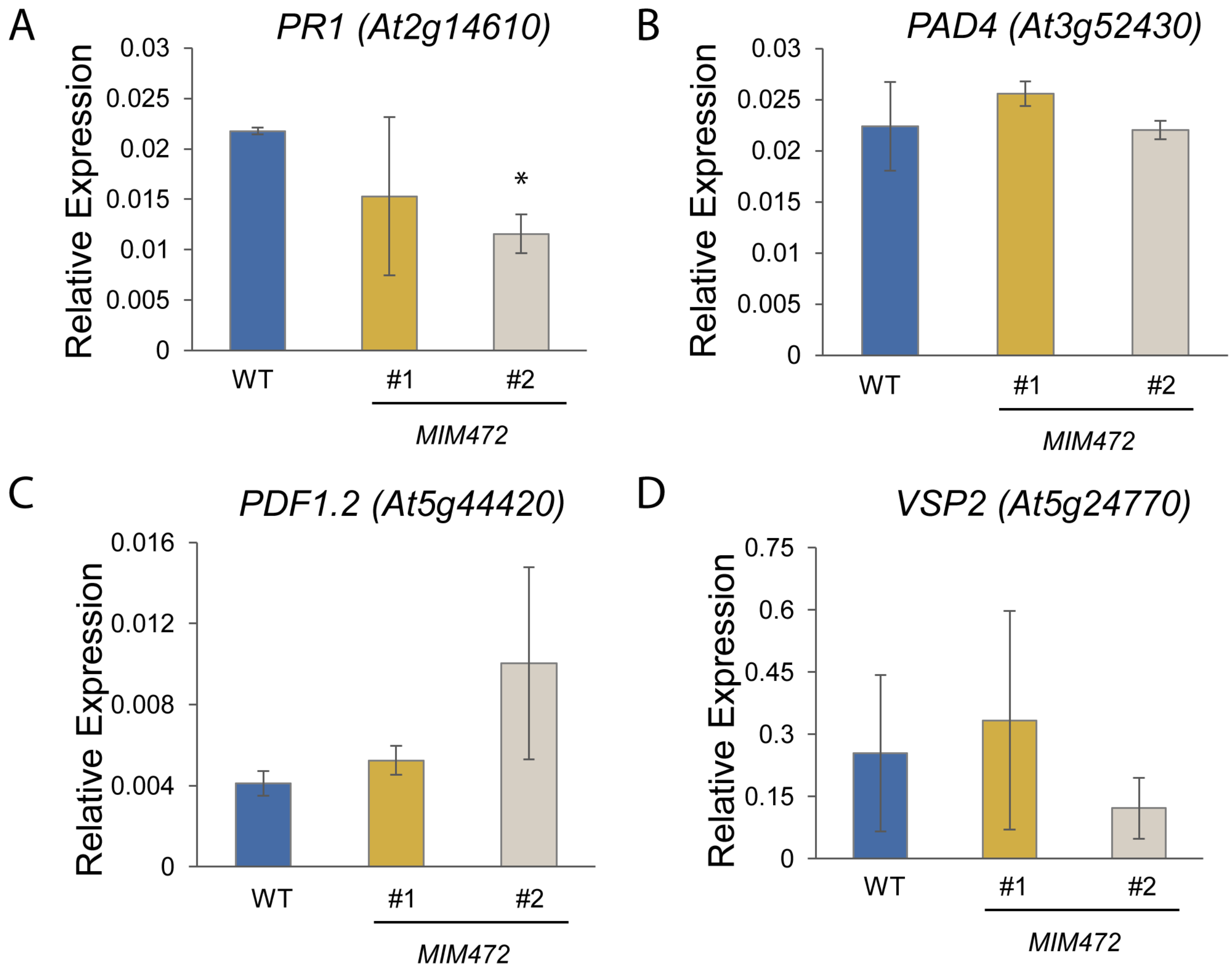
