## Supplementary material for "miR472 deficiency enhances *Arabidopsis thaliana* defence without reducing seed production": SOM Table 1

**Supplementary Table 1:** List of Oligonucleotides used in this study

| Primer name | Sequence (5’-3’) | Purpose |
| --- | --- | --- |
| AtmiR472_UPL RT | GTTGGCTCTGGTGCAGGGTCCGAGGTATTCGCACCAGAGCCAACGGTATG | cDNA synthesis |
| PB_AtmiR472_qPCR_Frw | GGCGGTTTTTCCTACTCCGCC | qRT-PCR |
| At5g43740_RTPCR_F | CAAGTCAATGGATGGCTTTCC | qRT-PCR |
| At5g43740_RTPCR_R | TTCATCAGGCTGCTCCACG | qRT-PCR |
| miR472 probe | GGTATGGGCGGAGTAGGAAAAA | miR472 blot |
| beta-tubulin2.s | GAGCCTTACAACGCTACTCTGTCTGTC | qRT-PCR |
| beta-tubulin2.as | ACACCAGACATAGTAGCAGAAATCAAG | qRT-PCR |
| PR1.QRT.s | GATGTGCCAAAGTGAGGTGTAA | qRT-PCR |
| PR1.QRT.as | GGCTTCTCGTTCACATAATTCC | qRT-PCR |
| PAD4.QRT.s | CGAATACATTGGTGACGAAGAA | qRT-PCR |
| PAD4.QRT.as | ACCCATTTTGCACTTGAACTCT | qRT-PCR |
| PDF1.2.QRT.s | CACCCTTATCTTCGCTGCTC | qRT-PCR |
| PDF1.2.QRT.as | GTTGCATGATCCATGTTTGG | qRT-PCR |
| VSP2.QRT.s | ACGGAACAGAGAAGACCGAC | qRT-PCR |
| VSP2.QRT.as | TCTTCCACAACTTCCAACGG | qRT-PCR |
